# Revealing heterogeneity of brain imaging phenotypes in Alzheimer’s disease based on unsupervised clustering of blood marker profiles

**DOI:** 10.1101/339614

**Authors:** Gerard Martí-Juan, Gerard Sanroma, Gemma Piella, for the Alzheimer’s Disease Neuroimaging Initiative and the Alzheimer’s Disease Metabolomics Consortium

## Abstract

Alzheimer’s disease (AD) affects millions of people and is a major rising problem in health care worldwide. Recent research suggests that AD could have different subtypes, presenting differences in how the disease develops. Characterizing those subtypes could be key to deepen the understanding of this complex disease. In this paper, we used a multivariate, non-supervised clustering method over blood-based markers to find subgroups of patients defined by distinctive blood marker profiles. Our analysis on ADNI database identified 4 possible subgroups, each with a different blood profile. More importantly, we show that subgroups with different profiles have a different relationship between brain phenotypes detected in magnetic resonance imaging and disease condition.

## Introduction

Alzheimer’s disease (AD) is the most common cause of dementia, a condition affecting more than 47 million people worldwide [1]. AD is one of the biggest concerns in global health care, due to its large economic and social impact. It is characterized by a deposition of amyloid-beta (Aβ) protein in the brain and the formation of tau plaques [2], and its most prevalent symptom is a progressive decline and deterioration of cognitive skills, leading to death. AD has been characterized as a multi-factorial disease [2, 3], involving many different processes and biological phenomena. Despite many efforts spent on research, we know relatively little about many aspects of the disease.

Currently, to diagnose accurately the disease and provide for an adequate treatment, several markers are used: cerebrospinal fluid (CSF) concentration of tau, p-tau and Aβ [4], and markers obtained from imaging techniques such as magnetic resonance imaging (MRI) to detect structural neurodegeneration, or positron emission tomography (PET) to detect Aβ concentration and tau deposition in the brain [5, 6]. Efforts have been made to find new, less-invasive markers in blood that can help diagnose the disease. In 2009, Schneider et al. [7] presented a review of proposed plasma marker candidates, concluding that a single reliable candidate had not been found yet. O’Bryant et al. [8], in their review on blood-based markers for AD detection, concluded that, although progress has been made, significant advancements on results validation are still needed before blood markers can be reliably used in clinical trials.

Several blood markers have been shown to correlate with structural changes. Dage et al. [9] studied the connection between neurodegeneration and tau protein levels in plasma, finding association between cortical thickness, cognition, and tau levels. In a study by Mattsson et al. [10], plasma neurofilament light was associated with cognitive deterioration and imaging markers of AD. Thambisetty et al. [11] analyzed the relationship between various plasma proteins, brain volume and cortical thickness in AD patients, finding links between plasma proteins and AD neurodegeneration. In a later work [12], they studied the relationship between plasma clusterin (apolipoprotein J) concentration and longitudinal brain atrophy, finding significant associations.

These previously described works do not take into account the possible heterogeneity arising from the interaction between blood and brain. AD is a highly heterogeneous disease, where its symptoms and path of degeneration can vary between patients, and identifying the different presentations or subtypes and their related signatures could help for a better detection of the disease and a better understanding of the interaction between different biological mechanisms (i.e. phenotypes). There have been many studies identifying possible heterogeneous subgroups of the disease. Noh et al. [13] found 3 subtypes on distinct patterns of cortical atrophy, using Ward’s clustering linkage method. Nettiksimmons et al. [14] presented subtypes based on CSF and MRI markers, and in a posterior study [15], they argued that vascular damage could explain subtyping difference. They also studied heterogeneity in mild cognitive impairment patients [16], using CSF, MRI, and plasma markers. In a different approach, Young et al. [17, 18] proposed an event ordering method to infer heterogeneous subtypes of the disease and stage.

Studies addressing heterogeneity of AD generally use the same data modality both to subtype the disease and analyze the obtained subgroups. This could hide relevant differences from other modalities. Moreover, clustering over features that are strongly correlated to the disease stage could lead to subgroups divided by disease severity, instead of by possible disease subtypes. Instead, we propose to use blood marker features to define the subgroups and then explore the relationships with the disease in each subgroup using brain volume and cortical thickness phenotypes and disease stages. We use a data-driven, non-supervised, multivariate clustering technique [19] to identify different presentations of the disease using blood markers, and then we analyze how the different blood profiles interact with brain structural phenotypes across the different disease stages. Compared to methods limited by univariate analysis, such as direct statistical tests for a single marker, multivariate analysis allows identification of potentially hidden blood marker profiles associated with latent pathological processes, addressing a limitation of the reported blood marker studies.

## Materials and methods

### Data

Data from the Alzheimer’s Disease Neuroimaging Initiative (ADNI) database (adni.loni.usc.edu) were used for this project. ADNI is a multi-site, longitudinal study launched in 2003 by Weiner et al. [20] that includes acquisitions of MRI, PET, other biological markers, and clinical and neuropsychological assessment tests of patients over time to track the pathology of AD. For this work, we have selected subjects that had available MRI T1.5 scan and blood marker information at baseline. After removing subjects that presented missing values, we ended up with a set of 298 subjects, including 52 cognitive normal (CN), 161 with mild cognitive impairment (MCI) and 85 with AD. Table 1 shows the demographic information of the studied cohort.

**Table 1.** Demographic information of the studied cohort.

|  | CN | MCI | AD | Total |
| --- | --- | --- | --- | --- |
| N<sup>o</sup> subjects | 52 | 161 | 85 | 298 |
| Age (years) | 75.1 ± 6.0 | 74.1 ± 7.5 | 74.6 ± 7.9 | 74.4 ± 7.4 |
| Sex (female) | 44.2% | 33.5% | 47% | 39.3% |
| Education (years) | 15.8 ± 2.7 | 15.8 ± 2.9 | 15.3 ± 3.0 | 15.7 ± 2.7 |
| ApoE4 | 9% | 52% | 67% | 48% |
| MMSE | 29 ± 1.2 | 26.9 ± 1.8 | 23.4 ± 1.9 | 26.3 ± 2.6 |

Categorical variables are expressed as counts and percentages. Continuous variables are expressed as mean ± standard deviation. CN: Cognitive normal. MCI: Mild cognitive impairment. AD: Alzheimer’s disease. ApoE4: Apolipoprotein E4. MMSE: mini-mental state examination.

We preselected 235 candidate plasma markers from the available cohort of biospecimen markers in ADNI, including 190 protein plasma markers gathered by the AD Metabolomics Consortium, plasma neurofilament light, Aβ proteins 40 and 42, and 41 aminoacids. For the final selection, we accounted for the reported quality of the samples and removed any marker that had missing values in any of the reported subjects. The final selection consisted of 172 markers. Experimental design and quality control methods are described in [31] (for the protein plasma markers) and in the ADNI website (for the rest of biomarkers). MRI data were processed and registered to a common space using Freesurfer’s *recon-all* [21]. We selected 39 volumes of structures from relevant subcortical regions of the brain defined in the default atlas of Freesurfer [22] and cortical thickness of the whole brain for interaction analysis. The full list of the selected plasma markers and brain regions can be found in supplementary files S2 and S3.

Each volume value was normalized by the estimated intracraneal volume of the subject. Both structural volumes and plasma markers were standardized to [0, 1] range before processing, We used min-max scaling, substracting the minimum value of each biomarker and dividing by the difference between the maximum and the minimum. This way, we preserve zero entries and introduce robustness to small standard deviations in the biomarkers.

## Methods

To find the different profiles, we cluster the patients using their blood markers, without using neither brain phenotypes nor diagnosis. We analyze the resulting clusters to find the blood profiles of each cluster. To find heterogeneous brain presentations in each cluster, we analyze the relationships between brain phenotypes in each cluster and disease stage, thus revealing the interactions between blood marker profiles and brain phenotypes across the disease stages. Fig 1 shows the pipeline of the method.

**Fig 1.**
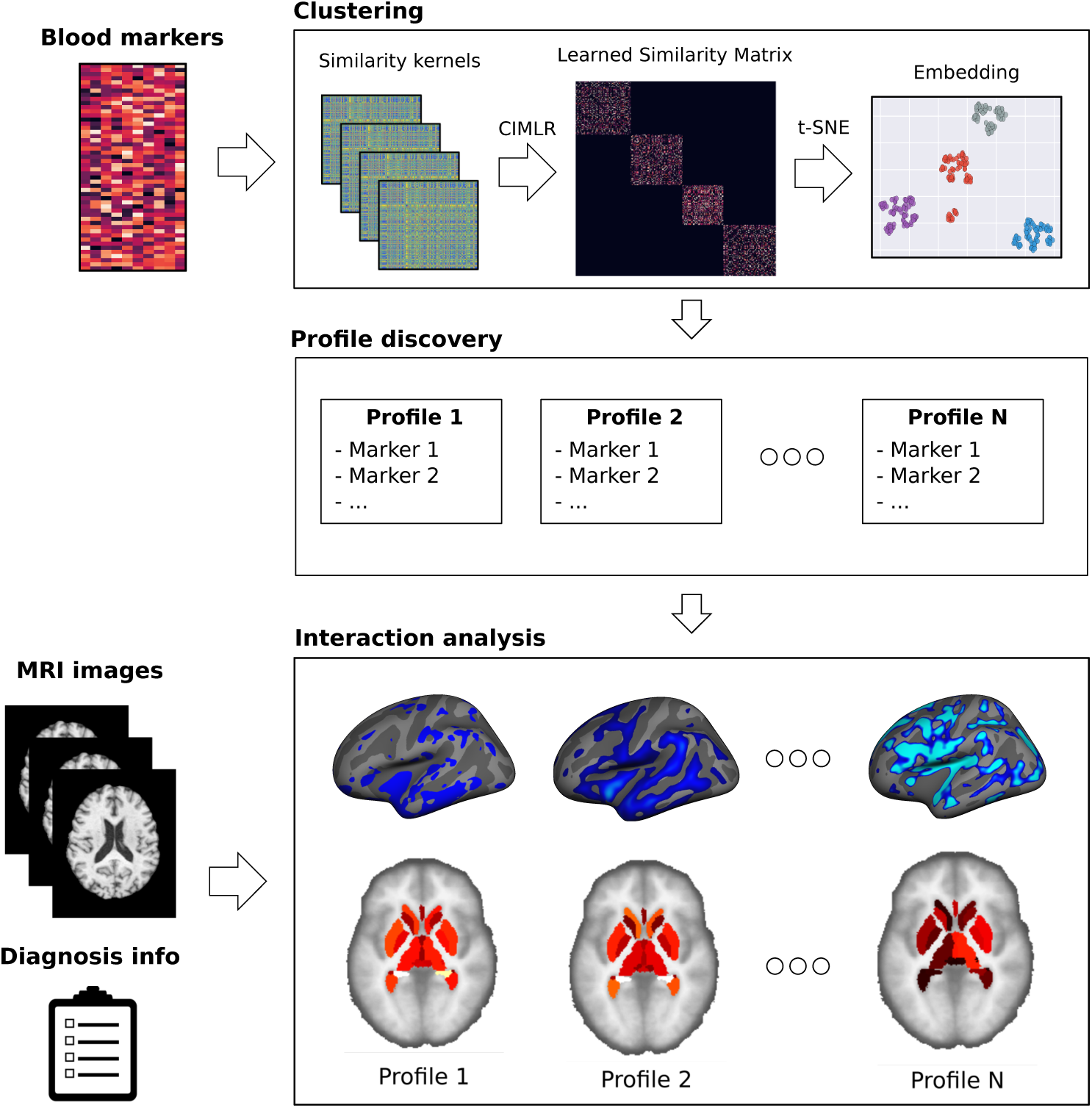
Method overview. We detect clusters on the space defined by the blood markers, define a different profile for each cluster, and, using the phenotypes extracted from the MRI images, we analyze the interactions between the profiles and the disease.

### Unsupervised clustering

We use CIMLR (which originally stands for cancer integration via multikernel learning, since it was developed for cancer subtyping) [19, 23], to identify the blood markers that reveal natural subgroups in the data, without taking into account neither the brain phenotypes nor the disease stage, to obtain subtypes not defined by disease stage. We could have used any other unsupervised clustering method, but we selected CIMLR, coupled with manifold learning and k-means clustering, due to its scalability with large amounts of data, good performance on a variety of datasets [19, 23], and interpretability of results.

CIMLR is a method based on multiple kernel learning that learns a similarity between each pair of samples by combining different kernels per feature (in our case, blood markers). It enforces a *C* block structure on the learned similarity, where each block is a set of samples similar to each other, i.e., a cluster. The number of clusters *C* must be specified beforehand. The learned similarity can then be used to compute a space of reduced complexity, where each subject is positioned with respect to the whole population, and the distance between subjects indicates how similar they are. By combining multiple kernels, each of which is based on a specific blood marker, it integrates the heterogeneous information, and provides the contribution of each blood marker in the computed low-dimensional representation.

Let the input data consist of *N* samples with *M* features (i.e., blood markers) each, be defined as 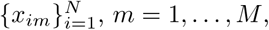 where *x_im_* represents the blood marker *m* of subject *i*. Each blood marker is assigned to *P* kernels *K_mp_*, *p* = 1*, …, P*. CIMLR solves for 3 variables: 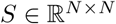 (the learned similarity matrix), 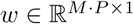 (a vector containing the weights associated to each kernel), and 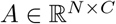 (a matrix enforcing *C* clusters in *S*). The optimization problem is defined as follows:

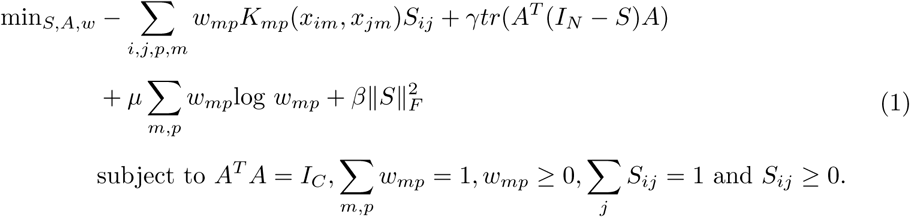

Here, γ*, µ*, and β are tuning parameters for the various terms of the optimization function, *I* represents the identity matrix, *||.||_F_* stands for the Frobenius norm, and *tr* denotes the trace of the matrix. The first term of Equation 1 links the learned similarity *S* with the combination of kernels from all features: similarity between two samples should be small if their kernel-based distance is large. The second term enforces *S* to have *C* connected components, through the auxiliary matrix *A* and its associated constraint *I_C_*. The third term imposes a constraint on *w* so that more than one kernel is selected, and last term applies a regularization penalizing the scale of the learned similarities. An extended overview of the algorithm, including the optimization method, can be found in [19]. A MATLAB implementation of the method by the authors of the paper has been used (https://github.com/BatzoglouLabSU/SIMLR).

We use Gaussian kernels to define *K_mp_*. In total, there are *P* kernels for each feature *m*, each with different parameters. This is needed because different markers could be sensitive to different ranges of parameters. We define *K_mp_* as:

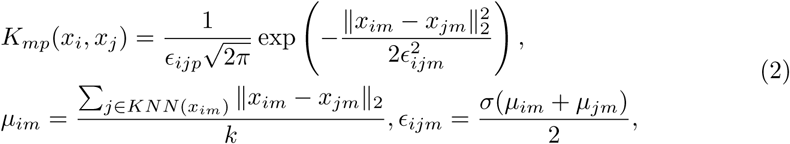

with *KN N* (*x_im_*) representing the *k* nearest neighbours of subject *i* with respect to marker *m*, and *||.||*_2_ being the Euclidean distance. For *k* ∈ {30, 45, 50} and σ ∈ {30, 35, 40, 45, 50}, we have a total of *P* = 15 kernels for each feature. The choice of parameters was done empirically. The method is mainly invariable to *P* [23]. For 172 markers, we optimize over a total of 15 × 172 = 2580 kernels.

As in [19], we estimate the best number of clusters with the heuristic proposed in [24], and further validate the choice with the elbow method. For visualization of the clusters and dimensionality reduction, we apply t-distributed stochastic neighbor embedding (t-SNE) [25], a manifold learning technique, on the resulting similarity matrix. After obtaining the low dimensional embedding, k-means clustering is used to discover the clusters.

### Cluster validation

We want to test the stability of the clusters against perturbations (e.g. particular choice of individuals). If the same clusters arise after modifying the choice of individuals, this suggests that the clustering is capturing an underlying structure in the data that is invariant to the particular choice of individuals, to some extent. We use a bootstrap procedure to test this stability. We apply the CIMLR-based clustering method to randomly select subsets of the data, with the same size of the original dataset but with replacement (i.e. some patients could appear several times, wheras others could not appear at all). After applying the clustering algorithm, we compute the similarity of the obtained clusters in each bootstrap iteration with the clusters obtained for the whole dataset.

We use the Jaccard index to quantify the similarity between clusters. This Jaccard index is defined, for two sets *A, B*, as the intersection divided by the union of the sets:

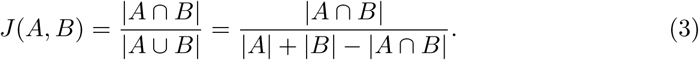

The higher the index, the higher the similarity and hence the stability.

### Profile discovery

Subjects in a given cluster share a specific profile of blood markers. To obtain the profile of each cluster, we need to determine which markers contributed more to the clustering. We look at the weights *w* in the optimization procedure: each weight accounts for the importance of a specific kernel. As described above, we generate 15 kernels for each of the 172 markers, with 15 associated weights. We compute the importance as the sum of those weights.

For further validation, we use one-way analysis of variance (ANOVA) tests of mean population across the clusters for the described markers to test whether the population of each subtype has a different mean. With this, we obtain a stable list of the most informative markers for the clustering, and describe the different blood profiles in each subgroup according to that set of markers.

### Interaction analysis

To detect different interactions between blood markers and structural brain phenotypes, we test for differences between the brain volumes and cortical thickness of the individuals in each subgroup. We perform three different comparison tests:

1. **Whole cluster analysis:** Each subgroup and the rest of the population, not taking into account diagnostic groups, to detect different characteristics in each subgroup.
2. **Diagnostic group analysis:** Diagnostic groups, for each subgroup, to find differences between stages of the disease in each subgroup with respect to the whole population.
3. **Diagnostic interaction analysis:** Each diagnostic group of each subgroup with the rest of subjects on the population in that diagnostic group, to detect different interactions between blood profiles and structural brain phenotypes across different stages of the disease.

We want to know whether cluster membership (independent variable) has significant effects on brain volume/cortical thickness (dependent variable). The significance thresholds may vary depending on the samples sizes, especially in our case that the groups may have different number of samples (i.e. clusters of different size). To correct for different sized groups and remove false positives, we used a permutation based method [26]. For a large enough number of iterations (1000), we perform random permutation of the independent variable (i.e. cluster membership), while preserving the cluster sizes and the diagnostic group sizes. Thus, we obtain a distribution of significance levels.

According to the permutation strategy, for the test to be significant, a higher significance level than those of most of the random permutations (e.g. 95%, 99%, … depending on the desired significance level) should be achieved. This correction is done to ensure that the obtained statistical significance was not caused by the different sample sizes of each subgroup and each diagnostic group. Fig 2 shows an outline of the procedure.

**Fig 2.**
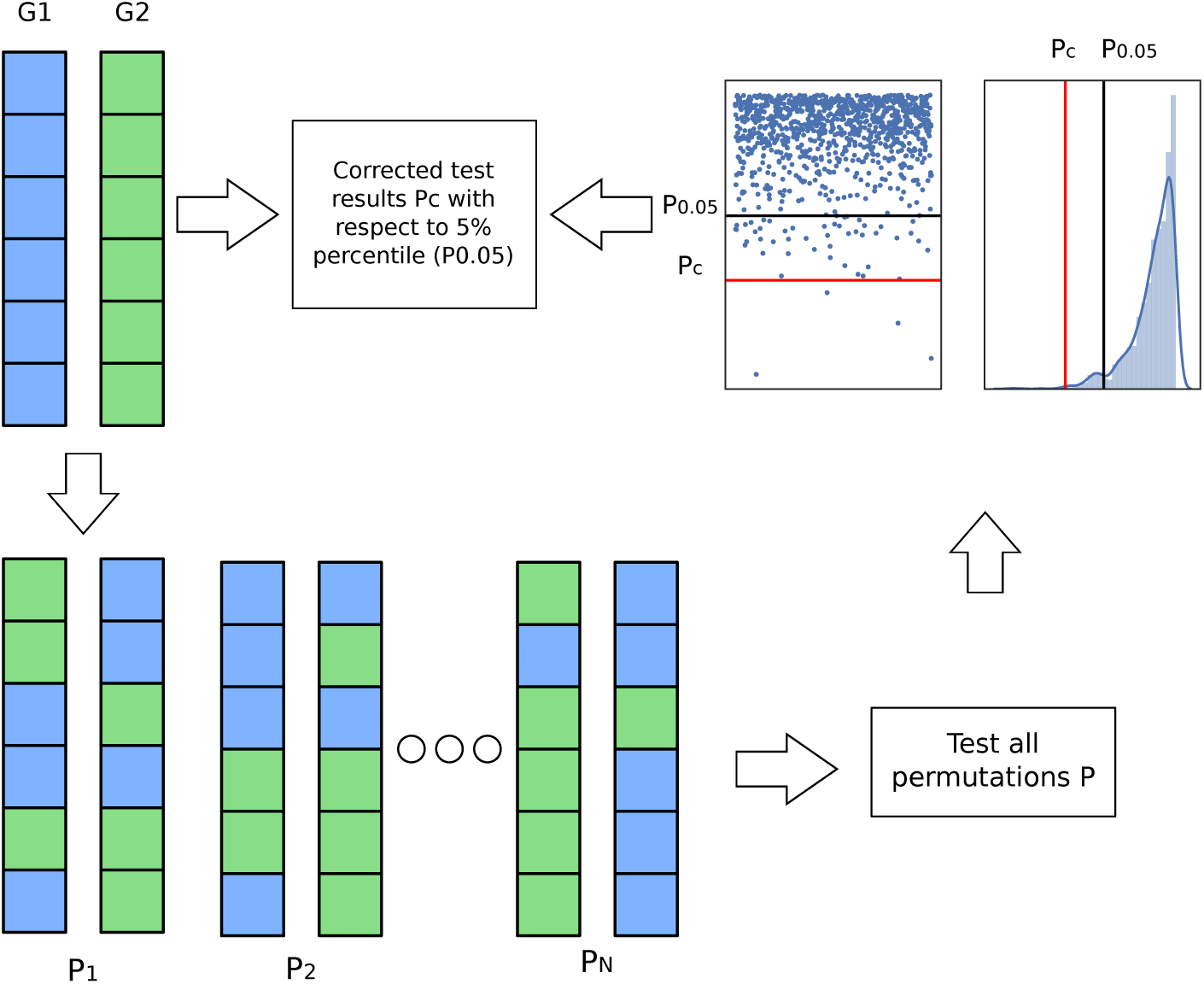
Permutation procedure. For groups G1 and G2, we create new random subgroups and test them. Then, we correct the obtained result in the original group (*P_c_*, red line), with respect to the 5% percentile of the obtained distribution of p-values (*P*_0.05_, black line). If the result is in the percentile, we consider it significant.

We use a non-parametric Mann-Whitney-Wilcoxon test for comparing the brain subvolumes. For cortical thickness, we use FreeSurfer’s *mri glmfit-sim* and *fsPalm* to implement the analysis. We also perform a cluster-wise correction on the surface of the cortex and applied Bonferroni correction for the two hemispheres. In this way, we can map corrected regions in the cortex that present significant differences in each analysis.

## Results

We applied the proposed method to the described cohort of subjects from ADNI database, to find the subgroups with heterogeneous blood profiles and analyze the interactions with the disease of each subgroup using volume features and cortical thickness. All the experiments are reproducible following the instructions found in the repository of the project https://github.com/GerardMJuan/simlr-ad.

### Clustering

After applying the heuristics on cluster size described in the Methods section, we obtained *C ∈ {*4, 6} using [24] and *C ∈ {*4, 5} using the elbow method. We decided on *C* = 4 as the most appropriate choice. Figure 3 shows the learned similarity matrix *S* and the cluster distribution in the first two dimensions of the low-dimensional space identified by t-SNE. We compared it to the similarity matrix and cluster distribution obtained using Euclidean distance on the original blood marker space. Fig 3 illustrates that clusters are not distinguishable when using an Euclidean-based similarity matrix, whereas CIMLR has a block structure that improves dimensionality reduction and cluster analysis.

**Fig 3.**
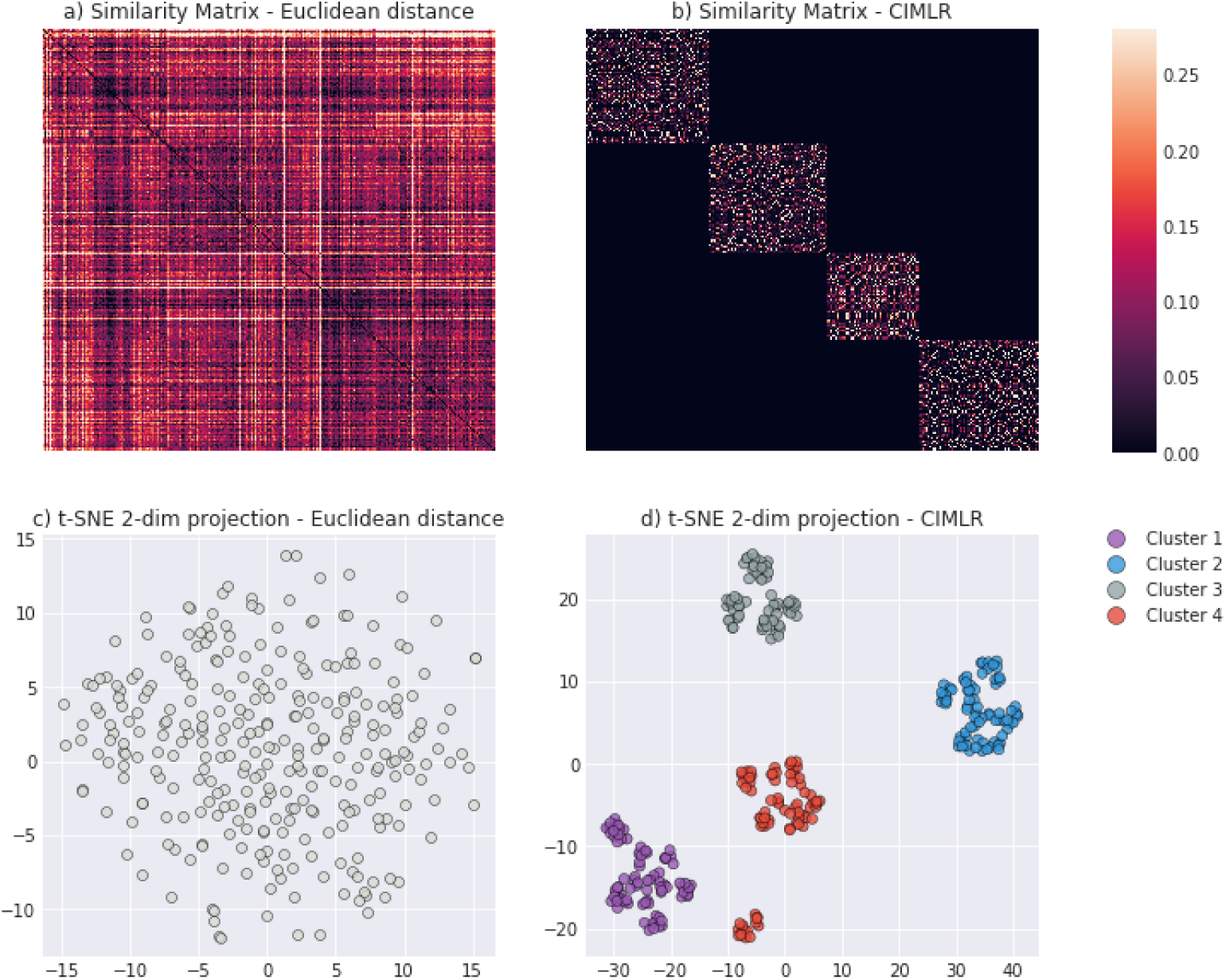
Similarity matrices and 2D embeddings. a) Euclidean distance between subjects over the initial blood marker space. b) Learnt similarity matrix *S*. Subjects in the matrix are ordered by the obtained clusters. c) d) 2D embeddings of the respective matrices in a) and b) using t-SNE.

We assessed the stability of the obtained clusters by using the bootstrap approach described in the Methods section and compared it with stability results obtained using a random clustering and k-means clustering with Euclidean metric. Table 2 shows the results. CIMLR got a larger mean similarity in each cluster than random clustering, and similar stability to k-means clustering. Clusters C1 and C2 appeared more stable than C3 and C4.

**Table 2.**
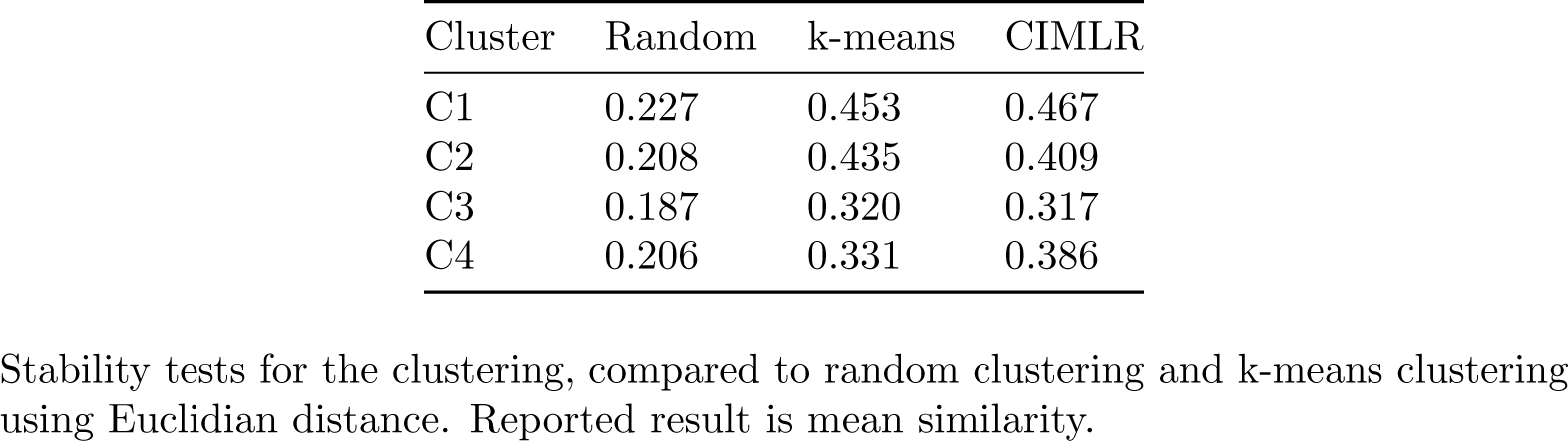
Stability tests.

Table 3 shows the demographic information of the subjects for each obtained cluster. Distribution of the diagnostic groups is similar to the distribution in the whole population, and the other characteristics (age, sex, education, Mini–Mental State Examination results and APoE4 genotype) are also similar across subgroups, with the exception of the mean age of C4, which is 5 years lower than the mean population, and a slightly higher fraction of women in C3 and C4. No major significant differences between subgroups were observed, meaning that the obtained groups were not biased.

**Table 3.** Demographic characteristics of the different clusters.

|  | Total | C1 | C2 | C3 | C4 |
| --- | --- | --- | --- | --- | --- |
| N° subj. | 298 | 82 | 77 | 61 | 78 |
| CN | 52 (17.4%) | 17 (20.7%) | 15 (24.5%) | 12 (19.7%) | 8 (10.2%) |
| MCI | 161 (54%) | 35 (45.4%) | 44 (57.1%) | 33 (54.1%) | 49 (62.8%) |
| AD | 85 (28.5%) | 30 (39%) | 18 (23.4%) | 16 (26.2%) | 21 (26.9%) |
| Age | 74.4 ± 7.4 | 77.2 ± 6.7 | 75.6 ± 6.1 | 75.1 ± 7.1 | 70.0 ± 7.2 |
| Sex | 39% | 25% | 28% | 52% | 53% |
| Education | 15.7 ± 3.0 | 15.9 ± 2.6 | 14.0 ± 3.4 | 15.6 ± 2.5 | 16.2 ± 3.0 |
| ApoE4 | 48% | 37% | 54% | 45% | 57% |
| MMSE | 26.3 ± 2.6 | 25.9 ± 2.9 | 26.6 ± 2.4 | 26.2 ± 2.7 | 26.5 ± 2.4 |

Categorical variables are expressed as counts and percentages. Continuous variables are expressed as mean ± standard deviation. Number next to the disease groups indicate the proportion of that group in the cluster. CN: Cognitive normal. MCI: Mild cognitive impairment. AD: Alzheimer’s disease. ApoE4: Apolipoprotein E4. MMSE: mini-mental state

### Blood marker ranking and profiling

CIMLR revealed patterns of plasma markers that relate to natural subgroups. Fig 4 shows the ten most relevant markers determined by the weight vector *w*. The method uses all blood markers to find the clusters but, unlike other methods, it automatically weights each marker. S4 data contains the full set of weights.

**Fig 4.**
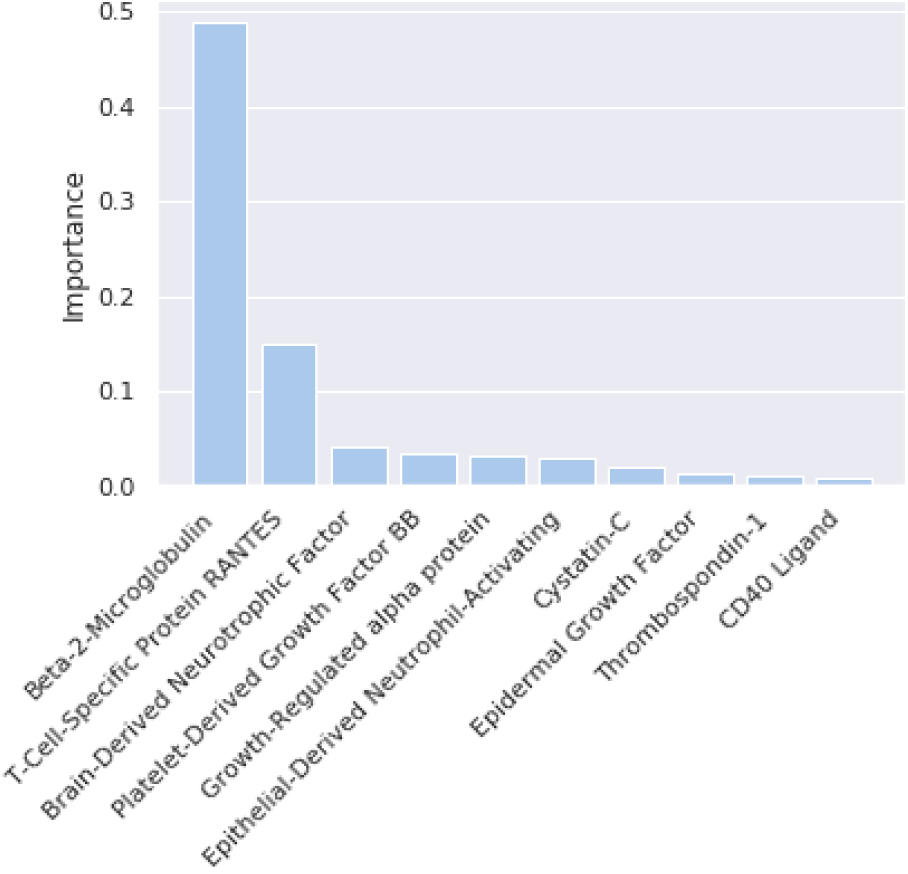
Marker weights. Top 10 marker weights in the kernel combination.

Fig 5 shows the values for each marker and cluster. All the ANOVA tests done for each of the described markers reject the null hypothesis with *p <* 0.001, meaning that the differences found between clusters on those blood markers are statistically significant.

**Fig 5.**
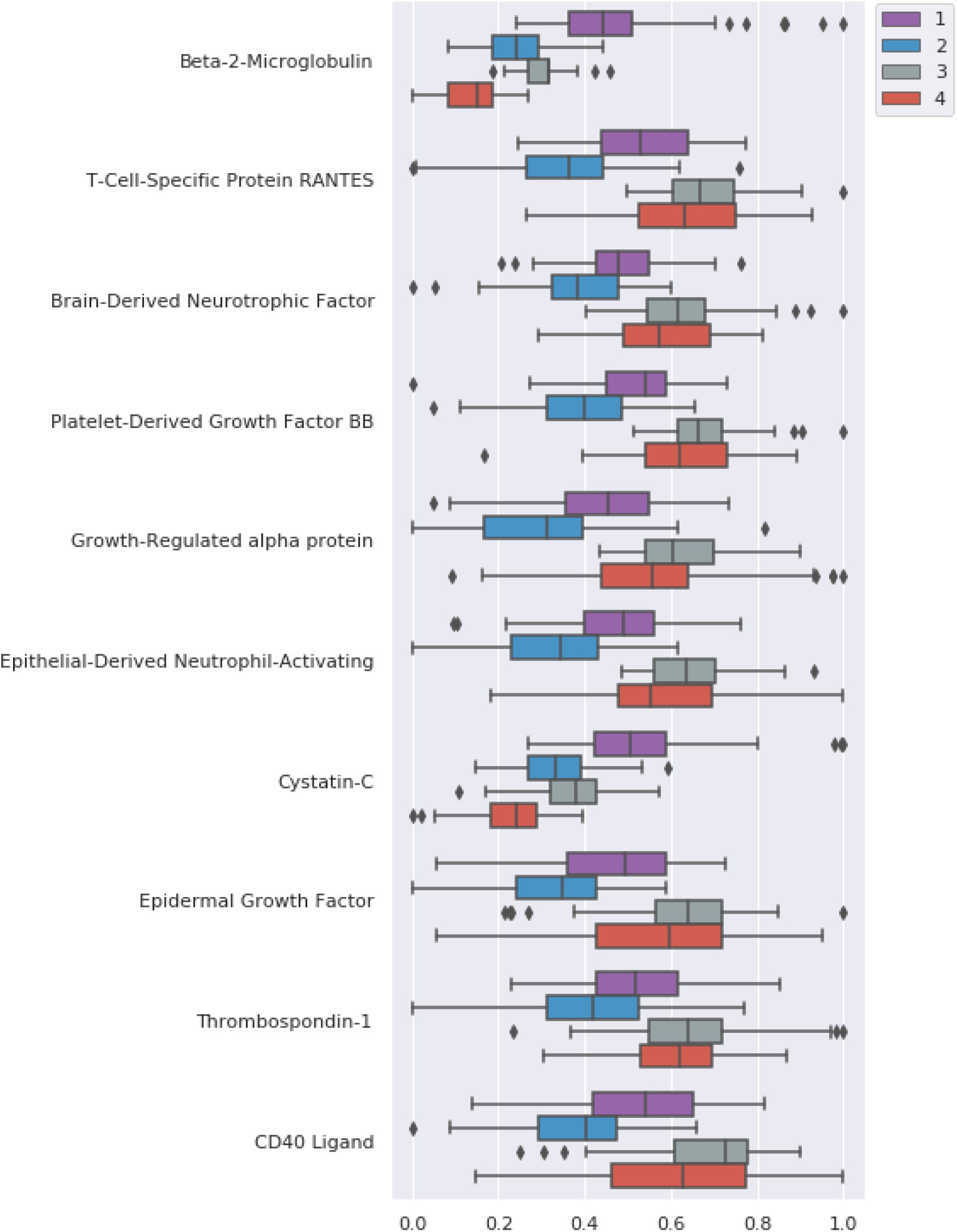
Marker distributions. Distribution of top ten ranked markers for each cluster. Normalized values.

Each of the profiles is defined by different values of the markers:

- C1 presents larger values of beta-2-microglobulin, cystatin-C and lower thrombospondin-1 with respect to the total population.
- C2 shows decreased values in all of the relevant markers compared to the whole population, with the exception of beta-2-microglobulin.
- C3 shows higher values in all of the relevant markers compared to the whole population, with the exception of beta-2-microglobulin and cystatin-C.
- C4 presents lower values of beta-2-microglobulin, cystatin-C and higher values of every other marker, compared with the general population.

Tau and amyloid-related markers, which are commonly associated with dementia [9, 27, 28], are not ranked highly. However, as per table 3, we know that the defined subgroups have a diagnostic distribution similar to the whole population: if markers that were highly correlated to the disease stage (such as the tau and amyloid related markers) had been selected by the algorithm, then that distribution would have been biased.

### Interaction analysis

We analyzed the heterogeneity between the different groups and the interactions between the stages of the disease and the brain volume and cortical thickness phenotypes depending on the blood profiles in each subgroup, as described in the Methods section.

### Whole cluster analysis

We compared subcortical brain volumes in each subgroup against the rest of the population, corrected for different group sizes and false positives using permutation tests. Fig 6 shows the differences in the characteristics of the population in each subgroup. C1 presents significant differences in the ventricles, putamen, accumbens area, left-vessel, right-pallidum, choroid plexus and posterior corpus callosum. C2 and C3 are similar to the general population: C2 only shows differences in the corpus callosum central and anterior, whereas C3 is only different to the rest in the right accumbens area. C4 shows many differences in the choroid plexus, ventricles, putamen and pallidum, among others.

**Fig 6.**
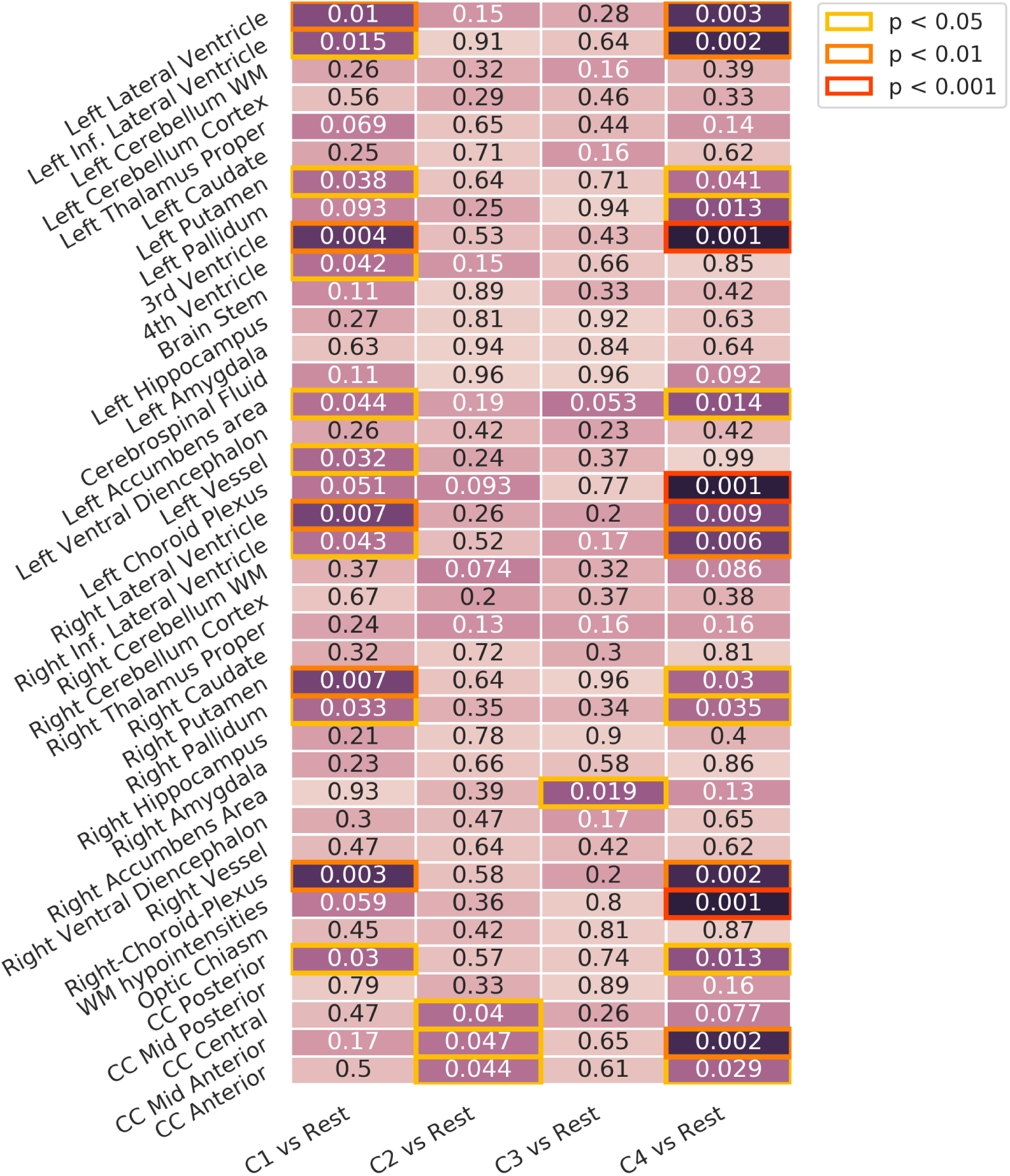
Whole cluster analysis. Differences in volume for each of the presentations against the rest. Corrected using permutation. Inf: Inferior. WM: White Matter. CC: Cingular Cortex.

We also tested for cortical thickness differences. Fig 7 shows the results. C1 presents differences in the superior parietal, supramarginal and central regions. C4 also shows differences in the supramarginal and central, and additionally in a region in the frontal cortex. We did not detect any differences in C2 and C3, which is consistent with the previous results on subcortical volume analysis.

**Fig 7.**
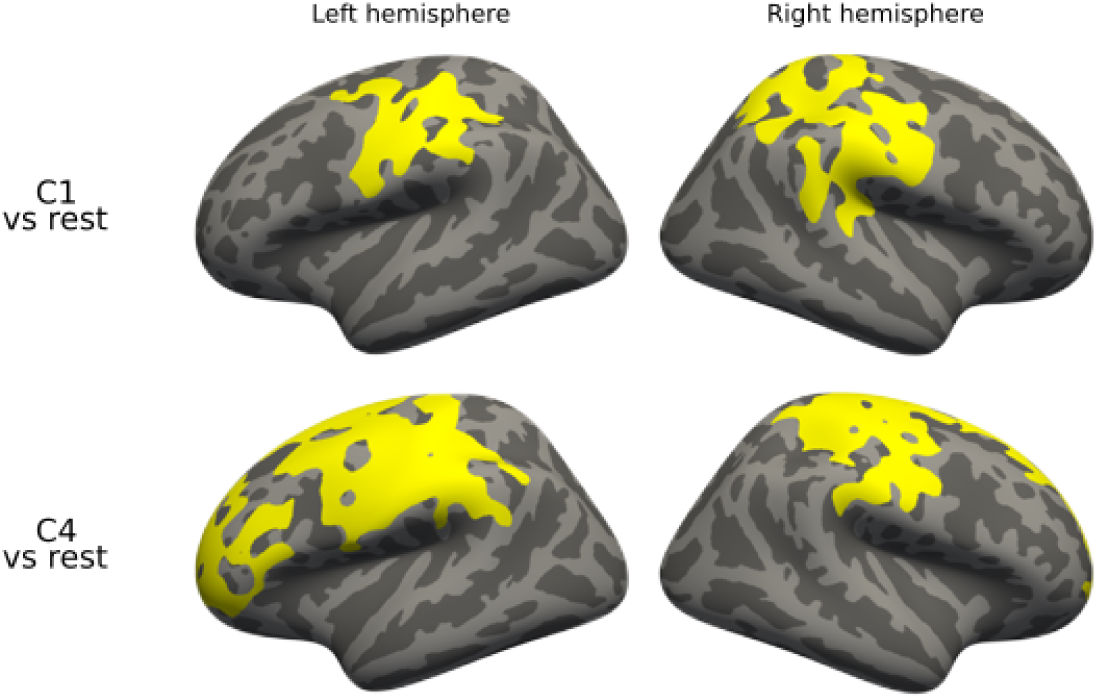
Whole cluster analysis, cortical thickness. Differences in cortical thickness for each of the presentations against the rest. Corrected using permutation.

### Diagnostic group analysis

To identify differences between diagnostic groups in each of the subgroups, we compared between diagnostic groups (CN, MCI, AD) for each of the different subgroups (C1 to C4). In this task, permutation tests allow us to detect differences between diagnostic groups that are specific to that subgroup, by correcting the result against random subgroups. Figs 8 and 9 show the difference between diagnostic groups in: (i) each of the subgroups (Fig 8) and (ii) the whole population (Fig 9).

**Fig 8.**
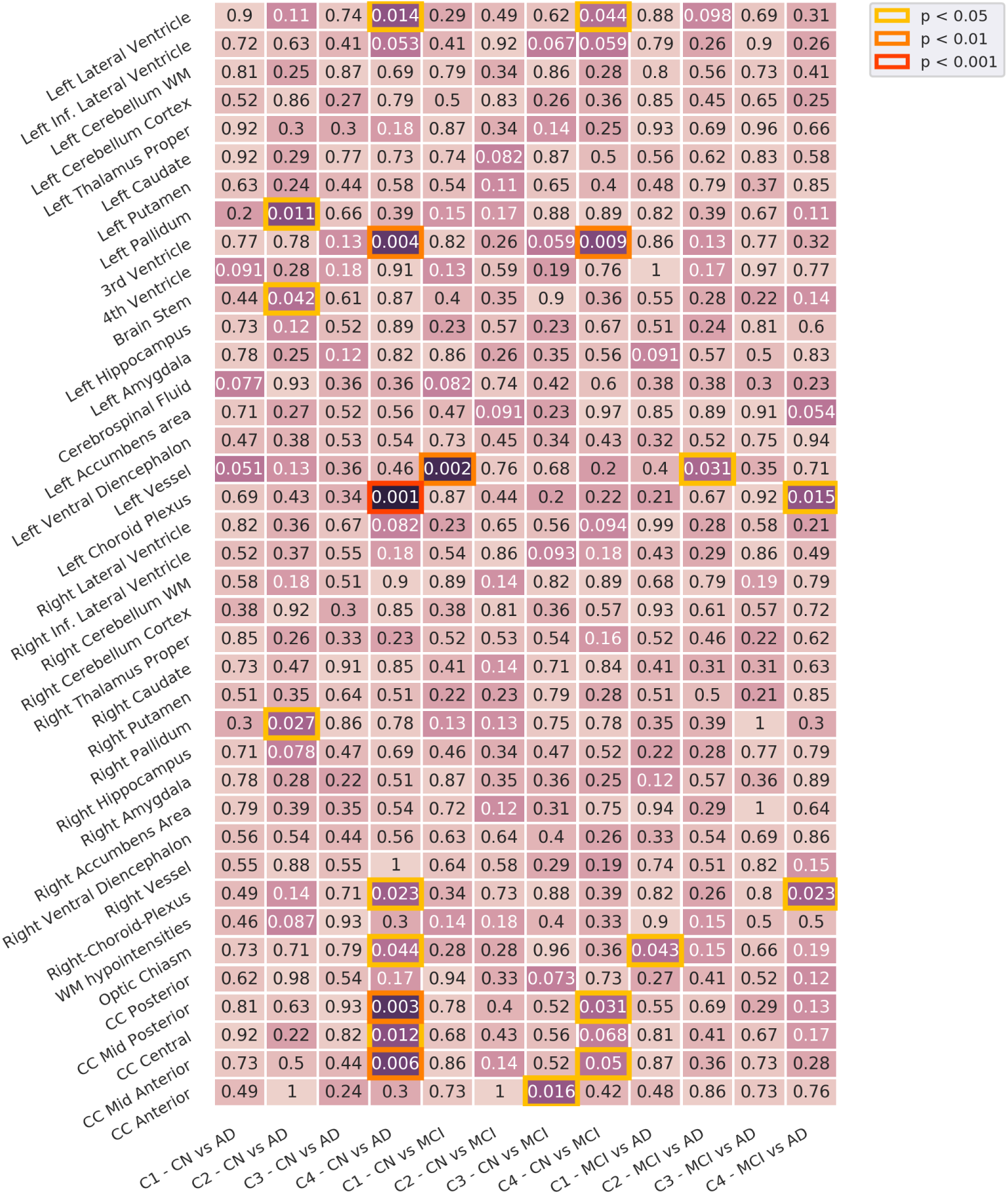
Diagnostic group analysis. Differences between diagnostic groups for each of the presentations. Corrected using permutation. Inf: Inferior. WM: White Matter. CC: Cingular Cortex.

**Fig 9.**
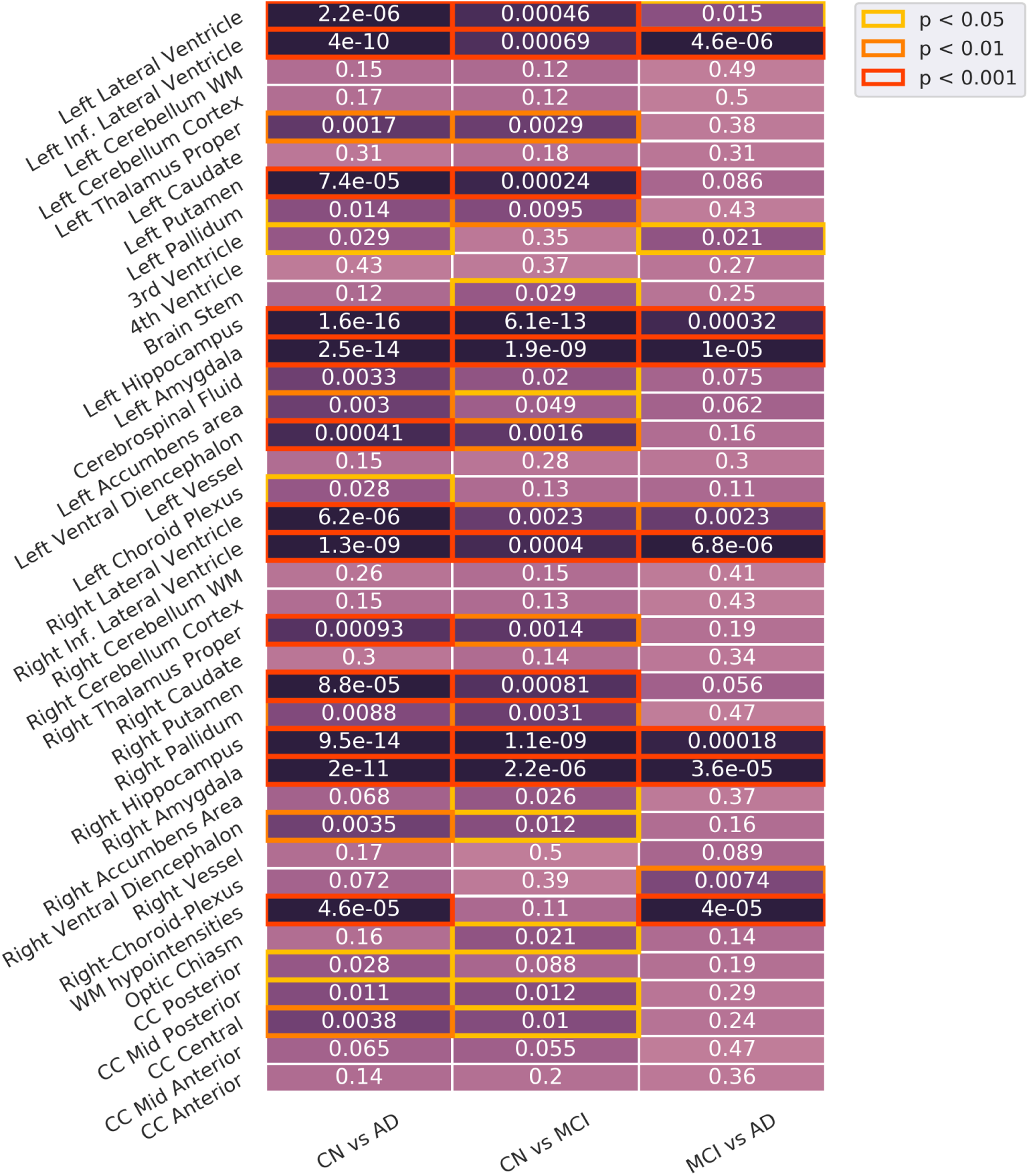
Diagnostic group analysis, whole population. Differences between diagnostic groups on the whole population. Inf: Inferior. WM: White Matter. CC: Cingular Cortex.

There are significant differences in C3 between CN and AD subjects in the corpus callosum, the third ventricle and the choroid plexus, and in C4 between CN and MCI subjects in the corpus callosum and the ventricles. C1 and C2 have more sparse differences with respect to the whole population. Most of the statistically significant differences correspond to volumes that show less significant differences on the whole population analysis. Intra-group heterogeneity between disease stages is located in specific regions that are usually less affected by the disease.

We only detected differences in cortical thickness when testing on CN vs MCI in C4, after correcting for multiple comparisons. Fig 10 shows the detected regions on the cortical surface, located on the frontal cortex and on the right temporal and parietal regions.

**Fig 10.**
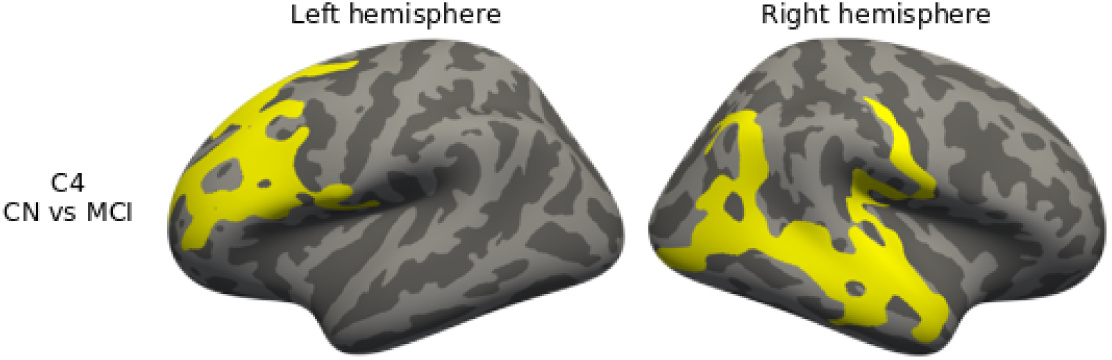
Diagnostic group analysis, cortical thickness. Differences in cortical thickness between diagnostic groups for each of the presentations. Corrected using permutation.

### Diagnostic interaction analysis

To detect different interactions between blood profiles and brain phenotypes across different stages of the disease, we tested for differences across same diagnostic subgroups for each cluster against the rest of the population with the same diagnosis. Fig 11 shows the results.

**Fig 11.**
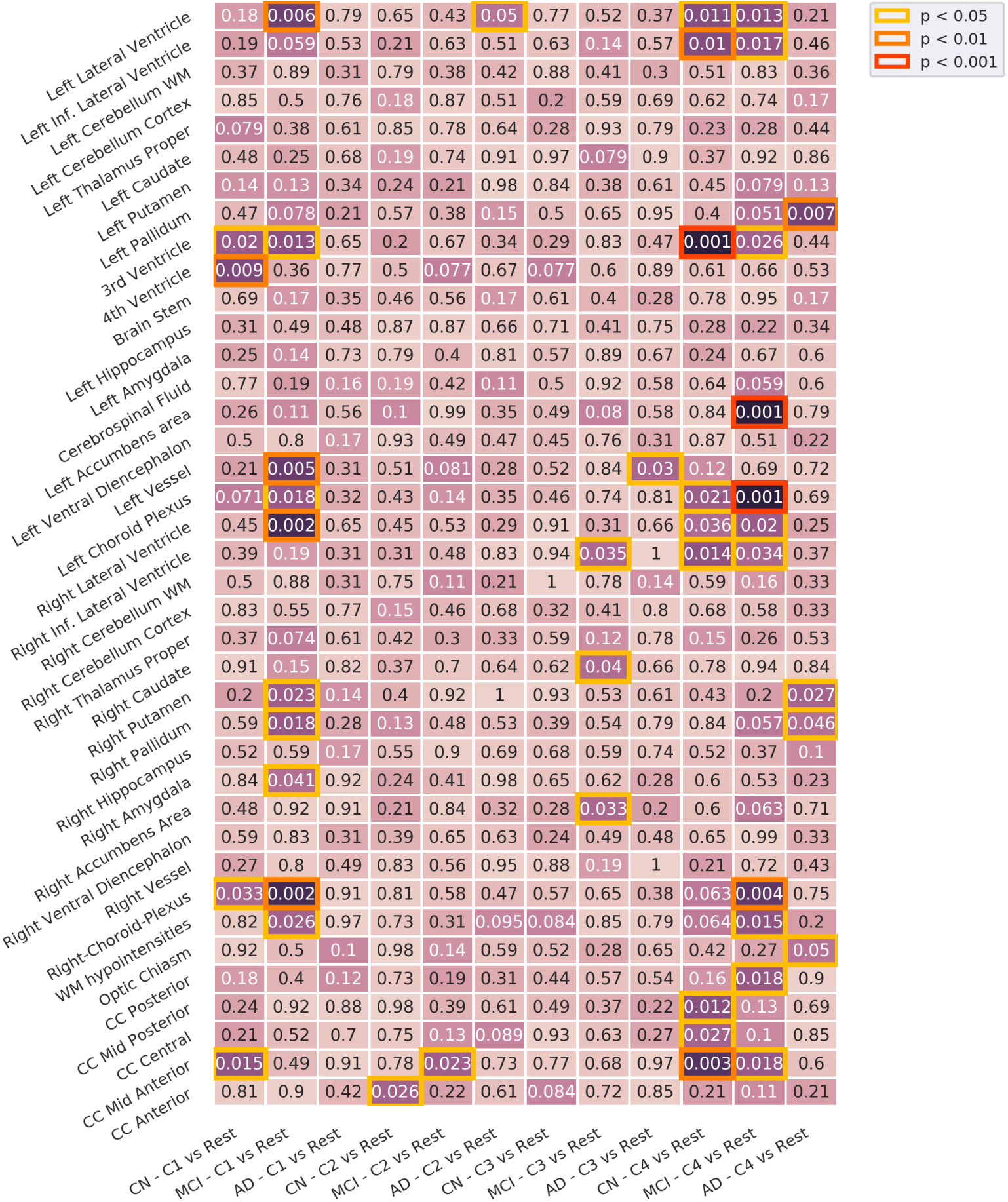
Diagnosis interaction analysis. Differences between diagnosis stages across presentations. Corrected using permutation. Inf: Inferior. WM: White Matter. CC: Cingular Cortex.

We observed differences in volume across all the subgroups:

- C1 differs in CN, presenting differences in the 3rd and 4th ventricles, and in MCI, with differences in the lateral ventricle, left vessel, left and right choroid plexus, among others. No differences were found in the AD group.
- C2 presents very small differences in the three diagnostic groups compared to the rest of the population.
- C3 does not present many significant differences: small volume differences in the MCI group, in the right accumbens, right caudate and right inferior lateral ventricle, and in the AD group in the left vessel.
- C4 shows many differences in the CN and MCI subgroups. CN presents differences in ventricles, both left and right, and the 3rd ventricle, and in various zones of the cingular cortex. MCI presents differences in the ventricles, left and right choroid plexus and left accumbens area, among others. AD also shows some differences esspecially in the left pallidum.

In cortical analysis, we found significant differences in MCI and AD patients of C4-which also presented many differences in our subvolume analysis-in the central and frontal regions of both hemispheres (see Fig 12).

**Fig 12.**
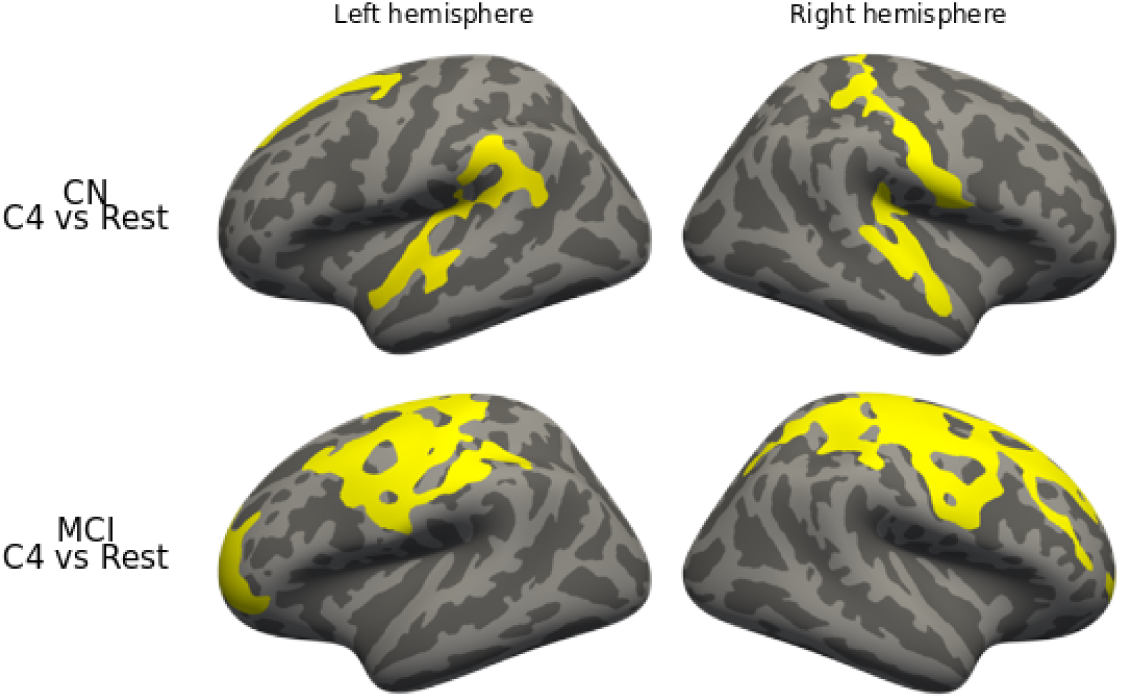
Diagnostic interaction analysis, cortical thickness. Differences in cortical thickness between diagnostic stages across presentations. Corrected using permutation.

## Discussion

We applied a multivariate data-driven procedure for AD subtyping using blood-based markers, to obtain heterogeneous groups with different blood marker profiles. We showed that patients with different profiles present different interactions between disease stage and brain phenotypes. Although existing blood markers can still not be used to properly diagnose the disease [7, 29, 30], using blood markers to detect patient profiles where the disease could behave differently and could provide valuable biological insights is a promising research direction.

The method identifies natural subgroups of patients in a multivariate way, finding hidden associations between the markers and automatically weighting the most important ones. The method is scalable to a large number of subjects and to a large number of features, allowing for an easier incorporation of other types of data, such as genotypes or other not-included blood markers. It also has some limitations: the obtained subgroups do not have a high stability. We also did not compare to other possible subtyping methods to further validate the obtained results.

From a clinical point of view, a study of the detected profiles is needed to investigate possible implications of the found markers. The different presentations of the disease detected in this work could be useful for a more personalized treatment in such an heterogeneous disease. Further validation of the results on a larger, independent cohorts of patients will be important to confirm the results and detect more complex profiles. Interactions of the profiles could also be further validated in other phenotypes, such as longitudinal brain atrophy.

## Supporting information

**S1 File. PTID list.** List of Patient ID (PTID) of the ADNI subjects used in this study.

**S2 File. Volumes List.** List of subcortical zones from FreeSurfer segmentation used in this study.

**S3 File. Blood markers list.** List of blood markers available in the ADNI website used in this study.

**S4 File. Full set of weights.** List of the weights assigned to each blood marker during computation.

## Acknowledgments

Data collection and sharing for this project was funded by the Alzheimer’s Disease Neuroimaging Initiative (ADNI) (National Institutes of Health Grant U01 AG024904) and DOD ADNI (Department of Defense award number W81XWH-12-2-0012). Data used in preparation of this article were obtained from the Alzheimer’s Disease Neuroimaging Initiative (ADNI) database (adni.loni.usc.edu) led by Principal Investigator Michael W. Weiner, MD, at UC San Francisco, and generated by the Alzheimer’s Disease Metabolomics Consortium (ADMC), lead by Dr. Kaddurah-Daouk, at Duke Medical Center.

